# 31° South: phenotypic flexibility in adaptive thermogenesis among conspecific populations of an arid-endemic bird - from organismal to cellular level

**DOI:** 10.1101/461871

**Authors:** Ângela M. Ribeiro, Clara Prats, Nicholas B. Pattinson, M. Thomas P. Gilbert, Ben Smit

## Abstract

In north-temperate small passerines, overwinter survival is associated with a reversibly increased maximum cold-induced metabolism (M_sum_). This strategy may incur increased energy consumption. Therefore, species inhabiting ecosystems characterized by cold winters and low productivity (i.e., low available energy) may be precluded from displaying an increase in maximum metabolic rates. To examine whether M_sum_ is a flexible phenotype in such challenging environments, and ultimately uncover its underpinning mechanisms, we studied an arid-endemic small bird (Karoo scrub-robin) whose range spans a primary productivity and minimum temperature gradient. We measured M_sum_, body condition, mass of thermogenic muscles and two indices of cellular aerobic capacity from populations living in three environmentally different regions. We found that M_sum_ was seasonally flexible, associated with aerobic capacity of limb muscles, but not increasing with lower temperatures, as predicted. Notwithstanding, the cold limit (temperature at which birds reached their maximum metabolic capacity) decreased in winter. These results indicate that birds from arid-zones may respond to cold conditions by altering thermosensation, rather than spending energy to produce heat in skeletal muscles.

## INTRODUCTION

Temperature influences animal life at all levels of organization. It affects the efficiency and functionality of biochemical networks and organism physiological responses (Hochachka and Somero 2002) in such a pervasive way that extreme heat or cold can be lethal. Therefore, to withstand thermal changes, vertebrates living in seasonal environments can reversibly alter their physiological phenotype in a process termed phenotypic flexibility (Angilletta *et al.* 2010) (McKechnie and Swanson 2010).

At cold temperatures, homeothermic-endotherm animals (i.e.: birds and mammals) loose heat from their warm bodies to the environment. To maintain a high body temperature in cold conditions, birds principally increase the rate of heat production through shivering (Hothola 2004). While shivering thermogenesis results from cellular processes, namely the activation of energetic metabolism pathways to burn cellular fuels to power skeletal muscle contraction (Hothola 2004), it can be quantified at the organism-level as the maximum thermoregulatory metabolic capacity (M_sum_; (Thompson 2010)). In fact, several studies showed that cold temperatures lead to increases in M_sum_ (Vezina et al. 2011) (Swanson et al. 2014a). One explanation for this flexibility in maximum metabolic capacity is the cold adaptation hypothesis, which posits that birds wintering in cold climates should have higher M_sum_ than those in warmer climates, and hence high M_sum_ is critical for overwinter survival in very cold regions (Marsh and Dawson 1989) (Swanson and Garland 2009). An alternative explanation is offered by the climate variability hypothesis (Janzen 1967), (Bozinovic and Naya 2014), which posits that broader climatic fluctuations results in wider flexibility in thermal tolerance as a means to cope with the fluctuating environmental conditions.

Several mechanisms, from whole-organism level down to the biochemical level, have been proposed to explain the high cold-induced M_sum_: i) increase in body condition assessed as body mass (Vézina *et al.* 2006) (Zheng et al. 2014); ii) increase in muscle mass, in particular the pectoralis muscle which is the primary thermogenic organ (Vézina *et al.* 2006) (Swanson et al. 2014a); iii) increase in enzymatic activity in oxidative metabolic pathways (Vézina *et al.* 2006) (Zheng *et al.* 2014) (Liknes and Swanson 2011a); and iv) increase in fat catabolism (Dawson et al. 1992) (Thompson 2010). However, evidence for flexibility in M_sum_, as well as support for the above mentioned mechanisms driving up-regulation of M_sum_, comes primarily from endotherms living in north-temperate ecosystems, where summer primary productivity is high (Prince and Goward 1995), and therefore individuals can afford the large energetic cost of up-regulating metabolic heat production (Hothola 2004).

In ecosystems characterized by low primary productivity and large seasonal temperature fluctuations, it is not clear how endotherms survive the low winter temperatures if not using energy saving strategies such as heterothermy. Therefore, to improve our understanding of the physiological underpinnings of adaptive thermogenesis in such ecosystems, we studied a small passerine (Karoo scrub-robin, *Cercotrichas coryphaeus*; hereafter designated scrub-robin) living in the subtropical arid-zone of southwestern Africa. Its range spans over an area of overall low primary productivity (Prince and Goward 1995) and exhibits a thermal gradient from west to east (Figure1C). As one transects from the Atlantic coast to inland, winters become increasingly colder, and primary productivity decreases as rainfall becomes increasingly unpredictable. Specifically, along the Atlantic coast winters are mild (mean minima: 8°C, record minima >0°C) and moderately productive from regular fog and predictable winter rainfall; in contrast, in the continental region minimum temperatures reach sub-zero (mean minima: -3°C, record minima <- 10°C) and primary productivity is very low year-round. Given these environmental features, the scrub-robin renders an ideal system for testing whether sub-tropical birds fine-tune phenotypes associated with adaptive thermogenesis in response to local environmental conditions. We hypothesize that birds from populations experiencing sub-zero winter temperatures would exhibit more pronounced increase of M_sum_ than populations from milder conditions, a pattern that can be driven by hypertrophy of pectoral (Swanson et al. 2014b) and lower limb (Isaac *et al.* 2014) muscles. However, low food abundance in such low productive ecosystems would hinder muscle build up. Thus, an alternative mechanism may be in place. Given the evidence showing that quantity and morphology of mitochondria (cellular powerhouses) change in response to bioenergetic cues (Putti *et al.* 2015) (Nasrallah and Horvath 2014), and that lipid droplets play an essential role in energy provisioning during exercise (Bosma 2016) and shivering in birds (Vaillancourt *et al.* 2005), we further predicted that the high metabolic capacity at organismal level would stem from increased cellular aerobic metabolism. Specifically, we anticipated that increased density of mitochondria and lipid droplets would be positively associated with M_sum_. To test our hypothesis we developed an integrative approach at several levels of organization (Figure1).

**Figure1.**
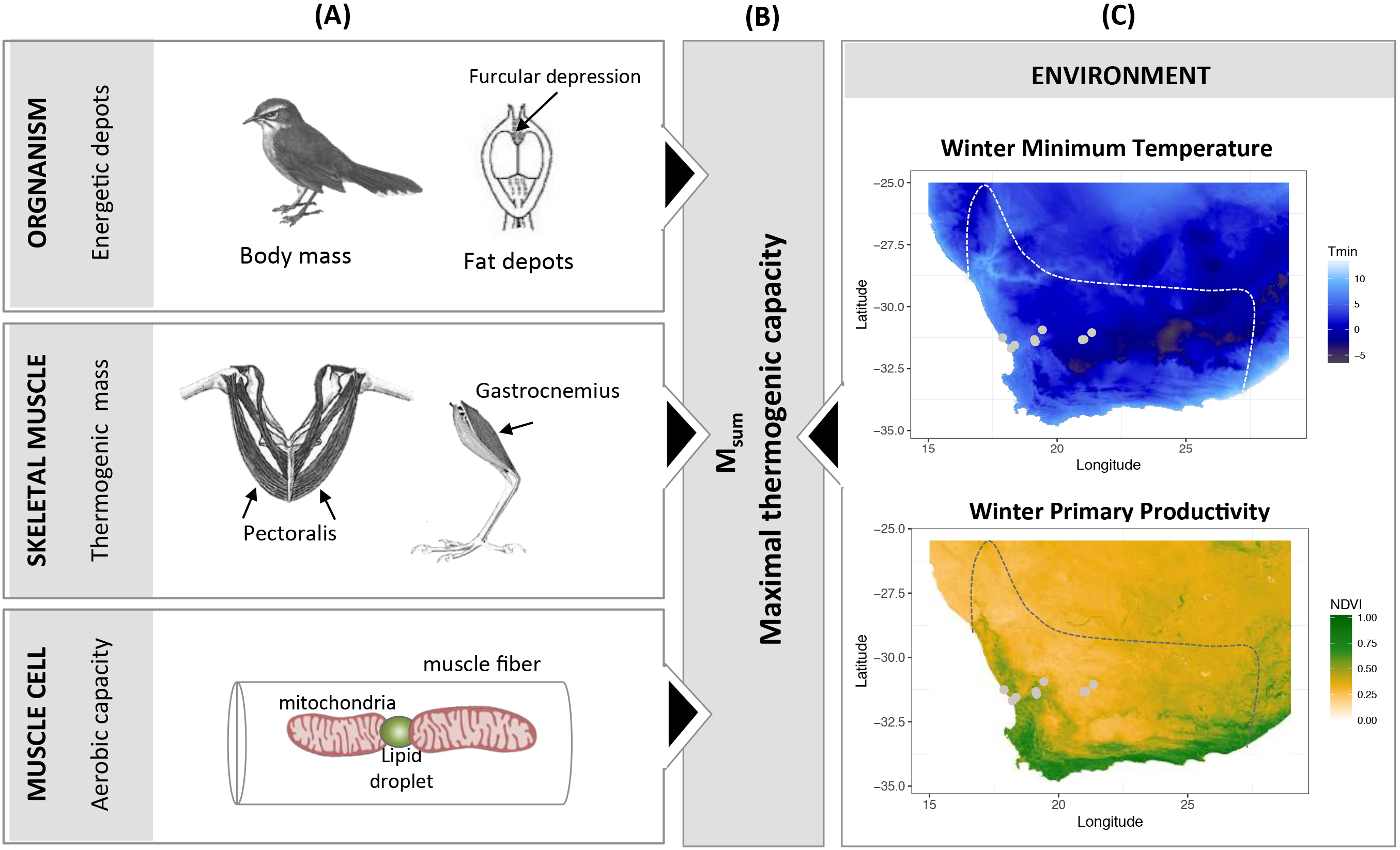
Framework to study the mechanisms of adap9ve thermogenesis in small birds living in arid-zones. (A) The underlying mechanisms driving (B) flexibility in maximal thermogenic capacity, and the role of (C) environmental variables in shaping those mechanisms. Study sites rela9ve to the species range (delimited by a dashed line) are depicted in the maps.

## METHODOLOGY

### Study sites, sampling and ethical clearance

We selected study sites that represent three main regions along the climatic and primary productivity gradient in the scrub-robin range (Figure 1C): *Coastal, Central* and *Inland.* We captured 85 adult birds in three regions and two seasons (summer: December 2015; winter: July 2016): n_coastal_ = 32 (n_summer_ = 18; n_winter_ = 14), n_central_ = 27 (n_summer_ = 14; n_winter_ = 13) and n_inland_ = 26 (n_summer_ = 10, n_winter_ = 16). Birds were captured using spring traps baited with mealworms (*Tenebrio molitor*). We kept them in individual cages in a quiet room with *ad libitum* access to food for no more than 24h, when the metabolic experiments took place. We released the birds at the point of capture, except those sacrificed (details below). Each sampling location was georeferenced and the GPS coordinates were the used to extract mean minimum temperature (T_min_) and Normalised Difference Vegetation Index (NDVI), a surrogate for primary productivity, for the months of data collection. T_min_ was extracted from two sources: i) WorldClim V2 database (http://worldclim.org/version2; (Fick and Hijmans 2017)) which is averaged from 1970-2000 interpolated at 2.5min resolution (~5km) and ii) South African Weather Service for three weather stations closest to our study and contemporary to our experiments. NDVI at 250m resolution from USGS-LandDAAC-MODIS dataset hosted by United States Geological Survey (https://lpdaac.usgs.gov/dataset_discovery/modis).

Permits to capture, handle and sacrifice birds were issued by the Northern Cape Department of Environmental Affairs (ODB 2665 & 2666/2015; Northern Cape Province) and CapeNature (0056-AAA008-00051; Western Cape Province) in South Africa. The Animal Ethical Committee at Nelson Mandela Metropolitan University (South Africa) approved all experiments (A15-SCI-ZOO-005).

### Body Condition: BM_scaled_, fat scores and mass of thermogenic muscles

The capacity to withstand environmental challenges (known as body condition; (Hill 2011) was assessed using three indices: i) body mass scaled by size (M_b-scaled_), ii) fat scores and iii) thermogenic muscles mass.

To quantify M_b-scaled_, which reflects the relative size of energy reserves such as protein and fat, we used the standardization technique proposed by (Peig and Green 2009): standard major axis regression between body mass (electronic scale, d = 0.01 g) and the linear body measurement tarsus-length (calliper, d = 0.01 mm). Thus, M_b-scaled_ = M_*i*_x [L_*0*_/L_*i*_]^bSMA^, where M_i_ and L_i_ are the body mass and tarsus-length of individual *i*, respectively; L_*0*_ is the tarsus-length arithmetic mean for the study populations to which index is standardized; b_SMA_ is the scaling exponent estimated by the standardized major axis regression of mass-length.

We checked fat accumulation at the furcular depression and abdomen in all 85 adult birds, and quantified it using a scale that ranges from zero (no fat) to eight (flight muscles not visible with fat covering the entire abdomen); (Kaiser 1993). Fat scores were verified in the birds we sacrificed to collect thermogenic muscles, as reported below.

We sacrificed 25 of the 85 birds: Coastal (n=10; 5 in each season), Central (n= 10; 5 in each season) and Inland (n= 6; summer: 1, winter: 5) by thoracic compression in the early morning after the rest-phase of birds. We excised the pectoralis and gastrocnemius muscles from the right-side body plane and measured their wet mass (electronic scale, d = 0.001 g). To estimate the total mass of pectoralis (Pectoralis_mass_) and gastrocnemius (Limb_mass_) we doubled the mass of the single muscles.

### Aerobic capacity: immunostaining and confocal imaging of mitochondria and lipid droplets

Because mitochondria density and morphology, and lipid droplets have been shown to play an essential role in energy provisioning during exercise, we measured the density of both intra-cellular organelles in the pectoralis and gastrocnemius muscles. Immediately after weighing the muscles, we fixed them by immersion into 2% paraformaldehyde as previously described (Dahl *et al.* 2014). In the laboratory, fiber bundles were teased apart under a stereomicroscope and stored until further processing. We prepare only muscles for birds at the extremes of the environmental gradient: *Coastal* (n = 10) and *Inland* (n = 6). Mitochondria were labelled by immunofluorescence using an antibody targeting Cytochrome C Oxidase and lipid droplets were stained using an antibody targeting Perilipin2. Briefly, single muscle fibers were permeabilized, incubated with primary anti-bodies, washed, subsequently incubated with secondary anti-bodies, and finally mounted in a glass slide. We used a Zeiss LSM710 microscope (Carl Zeiss, Germany) to image 8 - 10 fibers per individual muscle, and for each fiber we collected 16-22 z-planes. Orthogonal maximal projections were obtained for the 318 fibers, which were then used for image analysis.

To estimate the area occupied by mitochondria and lipid droplets, we developed a pipeline (step-by-step in Table S1, Supplementary Information) in CellProfiler Analyst (Jones *et al.* 2008). Our pipeline identified nuclei area, cytoplasm area, mitochondria area and lipid droplets area of each fiber. With these measures we estimated the fraction of the whole fiber occupied by mitochondria or lipid droplets as follows: Mitochondria fractional area (FAMito) = Mitochondria area /[cytoplasm area – nuclei area], Lipid droplets fractional area (FALipids) = Lipid droplets area /[cytoplasm area – nuclei area]. Transversal mitochondrial connections were counted manually from projected z-stacks. All processed images were manually curated by inspecting the overlay of the objects against the original image, and troubleshoot individually as needed. For full details on the immunostaining and image acquisition protocols see Supplementary Information.

### Cold-induced metabolism, body temperature and conductance

Birds showing any sign of body, wing and tail moult as well as those too agitated or not feeding enough (showing mass loss exceeding 5%) were excluded from cold-exposure experiments (n = 18). The remaining 67 birds were fitted with temperature-sensing passive integrated transponders (PIT-tag; BioThermo13, Biomark Inc., USA) to enable monitoring of body temperature (T_b_) throughout the experiments. Each PIT-tag was injected into the bird’s abdominal cavity as described in (Oswald *et al.* 2018).

After PIT-tag implantation, the bird rested in the cage for at least 30 min before cold-exposure experiments. Cold exposure experiments took place within 24h of capture at each field site, during the active phase of the birds (9am-4pm) after food was withheld for 2h.

To quantify metabolic rates under cold conditions, we used an open-flow respirometry system (FoxBox-C Field Gas Analysis System, Sable Systems, USA) to measure bird’s O_2_ consumption and CO_2_ production while exposed to a HelOx atmosphere (79% Helium + 21% Oxygen). HelOx was used because it allows maximum rates of heat loss at higher temperatures than normal air, and therefore prevents frostbite (Holloway and Geiser 2015). Data were recorded using EXPEDATA software (Sable Systems). Specifically, HelOx was pushed through a respirometry chamber (22 cm x 15 cm x 12 cm) holding the bird at flow rates of ~1.5 L min^-1^ using a mass flow controller (Omega, USA). The excurrent air from the respirometry chamber then passed through a multiplexer (Sable Systems), into an open manifold system, and was subsequently pulled through the gas analysers by the FoxBox system at a flow rate of around 0.5 L min^-1^.

The experiment involved placing the respirometry chamber housing the bird into a 40 L fridge/freezer (ARB, Australia) serving as an environmental chamber (Noakes et al. 2017). For the first 10-20 minutes of the trial, atmospheric air was pushed through the chamber (flow-rate of ~1.5 L min^-1^) to allow the bird to calm down. Trials started at air temperature raging from 15 – 20 °C (higher in summer than in winter) by switching the air stream to HelOx. The birds were then exposed to a sliding cold exposure protocol, by reducing air temperature by 3 °C every 10 minutes. Ambient temperature was recorded manually every 2 min using a Cu-Cn thermocouple (IT-18, Physitemp Instruments, USA) and temperature recorder (RDXL 12SD, Omega). Body temperature of birds was recorded at 1 sec intervals by placing the PIT-tag reader inside the environmental chamber. Trials ended when i) (Inline) started to decline indicating peak thermogenic metabolism had been reached, and ii) T_b_ < 34 °C. Unlike previously published studies on passerines undergoing cold exposure that were terminated when birds reached T_b_ < 37 °C (Swanson et al. 1996) we found that many scrub-robins showed a decline in T_b_ well below 37 °C, while still increasing resting metabolism and hence the reason for establishing the T_b_ < 34 °C threshold.

Cold exposure experiments generally took less than 1 h and peak metabolic rates were typically reached within 20 minutes of HelOx. At the end of each experiment the bird was placed back in the holding cage, in a warm place, with food and water available ad libitum. For each bird, we recorded the time, air temperature and T_b_ at which M_sum_ was reached. The air temperature at which M_sum_ was reached is defined here as the cold limit (Tc_L_). To calculate M_sum_, we obtained the highest 5 min mean (Inline) during cold exposure. We chose (Inline) over O_2_ consumption, as our field set up did not allow for systematic drift correction in oxygen values in all individuals. In addition, we estimated thermal conductance by calculating the rate of heat loss when birds reach their M_sum_ using the following equation C = M_sum_/(T_b_-T_a_), where T_b_ and T_a_ represent body temperature and HelOx temperature measured within five minutes of the birds reaching M_sum_, respectively.

### Statistics

We extracted NDVI and T_min_ values for our sampling sites using the R *Raster* package (Hijmans 2017). For all variables, we tested for normality and homogeneity of variance using Shapiro– Wilk’s and Levene’s test, respectively. If heteroscedasticity was detected, the response variable was log-transformed. We used an analysis of variance to test for sexual dimorphism. In the event that sex was not significant, we removed the variable and proceeded with pooled sexes. We tested for the role of environmental features such as T_min_ and NDVI on body condition, mass of thermogenic muscles (mass of pectoralis and gastrocnemius) and cellular aerobic capacity (density of mitochondria and lipid droplets), cold limit (T_CL_) and thermal conductance using generalised linear models (GLMs). We opted to use T_min_ experienced during each study period as a linear predictor instead of “season” as a categorical predictor as we believe that variation amongst study site in the former metric would explain physiological responses better.

Additionally, we examined the influence of T_min_ and NDVI, body condition, size of thermogenic muscles and cellular aerobic capacity on M_sum_ using GLMs.

All statistical analyses were performed in R v3.3.2 (R Development Core Team) and plots produced with ggplot2 package (Wickham 2016). We accepted p ≤ 0.05 as a significant difference for all statistical tests.

## RESULTS

Regardless of the source of T_min_ (WorldClim or SAWS) the results were consistent; therefore for the sake of brevity, in the main text we present the results for T_min_ obtained from WorldClim and report results with T_min SAWS_ in Supplementary Information.

### Body condition: BM_scaled_, fat depots and muscle mass

We found that minimum temperature (T_min_), but not primary productivity (NDVI), was a significant predictor of size corrected body mass (M_b-scaled_). Birds significantly increased body condition as T_min_ decreased (GLM; t_Tmin_ = -3.666, p < 0.01); FIGURE 2A-B). At the regional level, the decrease of T_min_ and NDVI lead to a significant increase of M_b-scaled_ for birds living in the *Inland* (GLM, t_Tmin_ = 2.254 p=0.03; t_NDVI_ = 3.229, p < 0.01) and *Coastal* (GLM, t_Tmin_ = -2.763, p = 0.01; t_NDVI_= -2.506, p = 0.02) regions, although no change was observed for *Central* region birds.

**Figure2.**
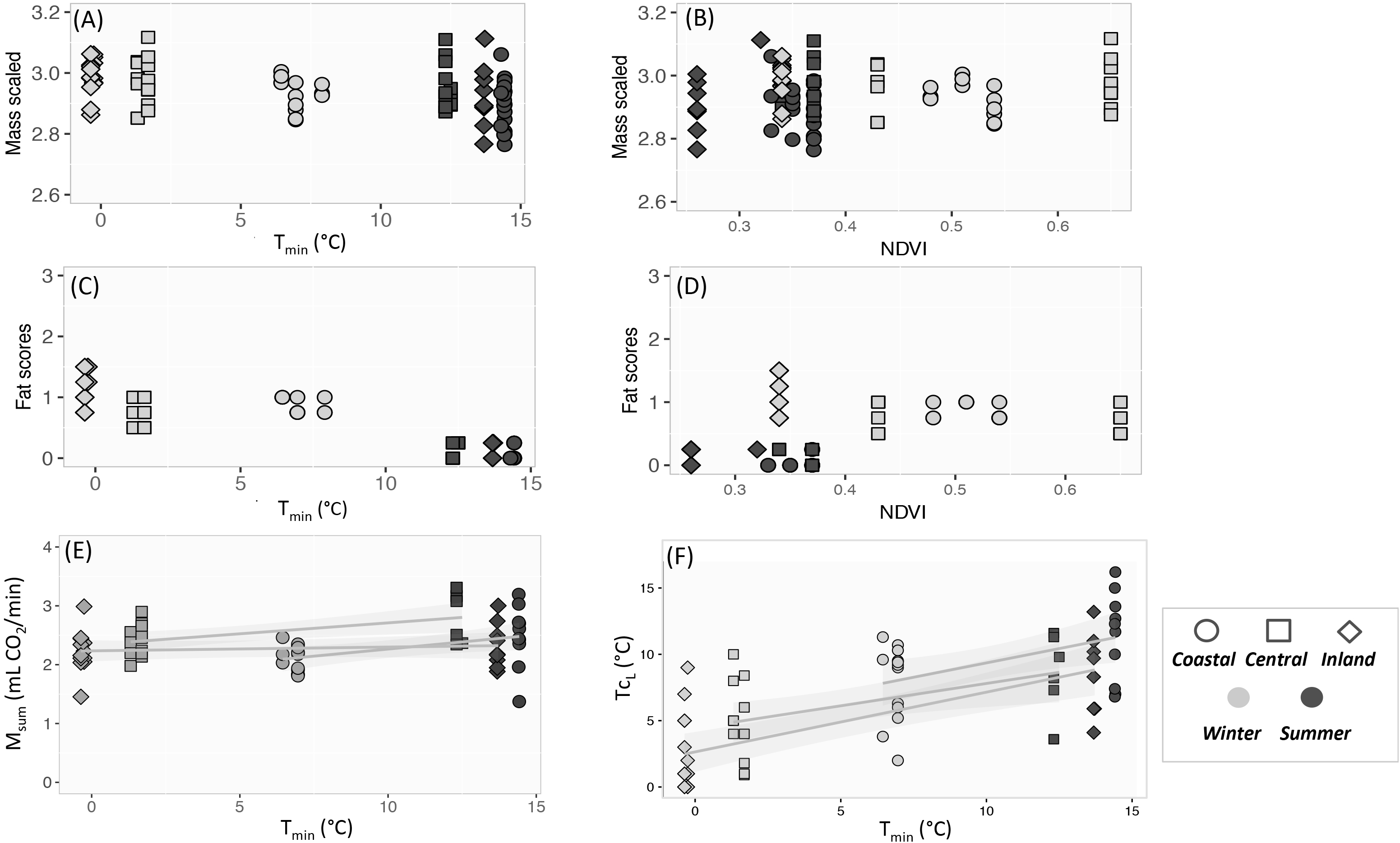
Varia9on in physiological traits in response to mean minimum temperature (T_min_) and primary produc9vity (NDVI). (A-B) Mass scaled, (C-D) Fat scores, (E) maximum thermogenic metabolic capacity (M_sum_) and (F) cold limit (T_cL_).

Minimum temperature, but not NDVI, was significantly associated with fat deposits (GLM, t_Tmin_= -14.795, p < 0.01; FIGURE 2C-D). The amount of visible fat significantly increased as T_min_ decreased in *Coastal* (GLM, t_Tmin_= -4.295, p < 0.01), *Central* (GLM, t_Tmin_= -6.245, p < 0.01) and *Inland* (GLM, t_Tmin_ = -2.421, p = 0.02) regions. For all populations fat depots were absent to very low during the summer (Fat scores = 0 - 0.25) compared to winter values ranging from 0.5 to 1.5 (1.5 = furcular depression almost completely covered with fat, plus small stripes of fat in abdomen).

Overall, the pectoral muscles (n = 15, mean ± SE = 1.114 ± 0.034g x 2 = 2.228 ± 0.068 g) accounted for 12.3% of scrub-robin body mass (n = 15, mean ± SE: 18.8g ± 1.382 g), while the gastrocnemius (n = 15, mean ± SE: 0.089 ± 0.002 g x 2 = 0.178 ± 0.004 g) represented 0.94% of birds’ body mass. There was no association of Pectoral_mass_ or Limb_mass_ with T_min_ or NDVI: GLM_pectoral_, t_Tmin_ = -0.832, t_NDVI_ = -1.116, p > 0.10; GLM_limb_, t_Tmin_ = -0.832, t_NDVI_ = -1.116, p > 0.10.

### Aerobic capacity: Mitochondria and lipid droplets density

Confocal micrographs of the pectoralis muscle revealed densely packed round mitochondria (Figure3A). Mitochondria fractional area (FAMito) ranged from 0.615 (± 0.084) in pectoral fibers, to 0.416 (± 0.092) in limb (Table S2, Supplementary Information). FAMito_pectoral_ was approx. 20% significantly larger than FAMito_limb_ (F = 38.054, p < 0.01).

**Figure3.**
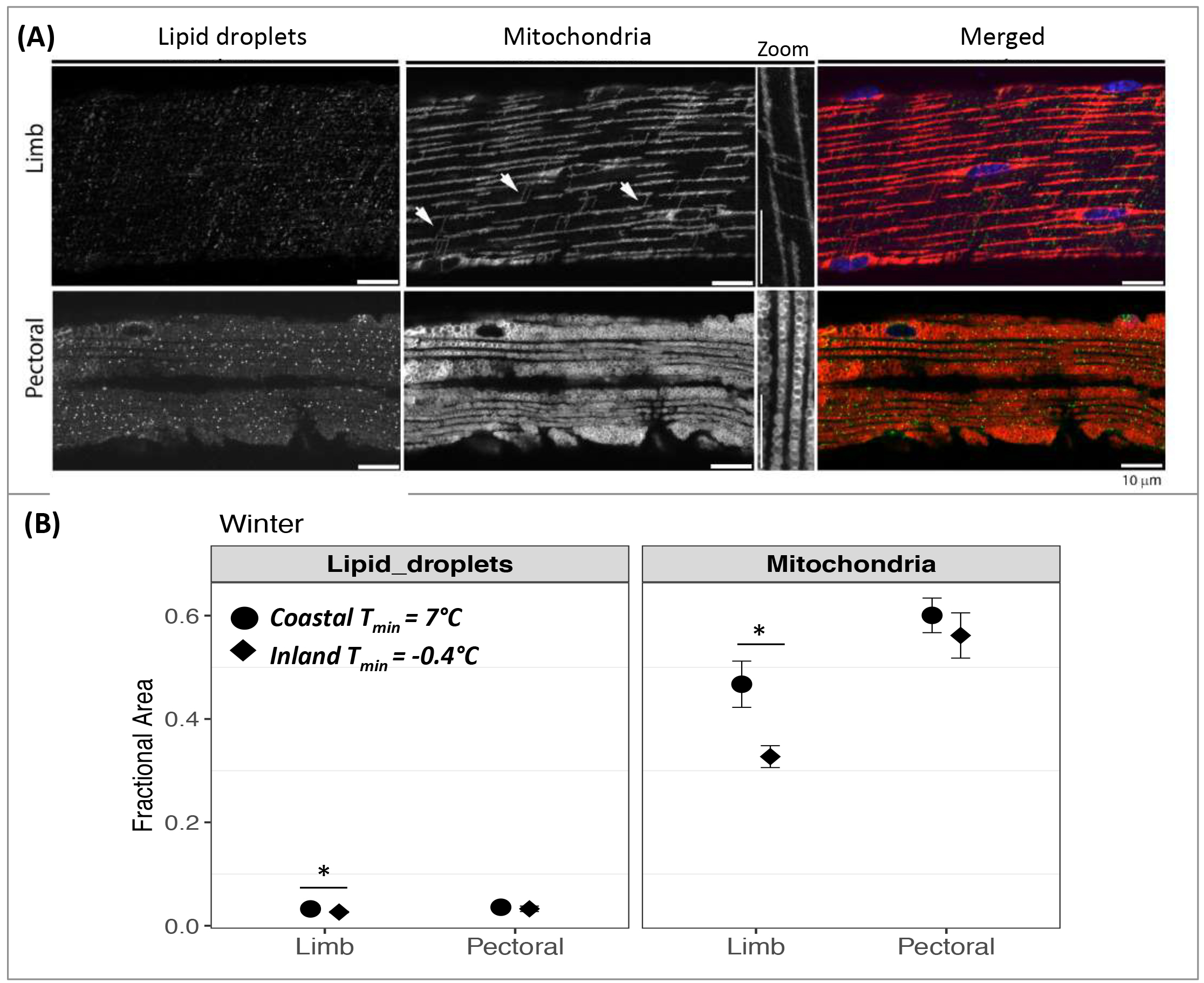
Variation in aerobic capacity in thermogenic muscles. (A) Confocal micrographs depicting mitochondria and lipid droplets (Perilipin2). Merged image show mitochondria (red), lipid droplets (green) and nuclei (blue). Transversal connections between mitochondria, exclusive of limb fibers, are indicated with arrows and visualized in zooming in images. Bars: 10 μm. (B) Portion of muscle fiber occupied by lipid droplets and mitochondria in limb and pectoral, in winter, in *Coastal* and *Inland* populations. Significant differences (α = 0.05) are annotated with an asterisk.

FAMito_pectoral_ was not associated with T_min_ (GLM, t = 2.071, p = 0.06) or NDVI (GLM, t = - 0.287, p > 0.10). FAMito_limb_ was not associated with NDVI (GLM, t = 1.803, p= 0.096) but associated with T_min_ (GLM, t = 2.994, p = 0.01): mitochondria densities increased with T_min_. Transversal connections between intermyofibrillar mitochondrial networks were exclusively observed in limb fibers (Figure3A). In the *inland* population, 80% of the individuals showed connections, while in the *coastal* population connections were only present in 40% of the birds.

The mean fractional area occupied by lipid droplets (FALipids) ranged from 0.035 in pectoral fibers to 0.040 in limb (Table S2, Supplementary Information), with no significant difference between them (F = 0.195, p = 0.66). FALipid_limb_ was significantly associated with T_min,_ but not with NDVI (GLM, t_Tmin_ = 1.794, p = 0.04; t_NDVI_ = -0.617 p > 0.1), showing an increase at higher temperatures (Figure3B). FALipid_pectoral_ did not change with T_min_ or NDVI (GLM, t_Tmin_ = 1.082, t_NDVI_ = 0.486, p > 0.1).

### M_sum_ association with environmental features, body condition, thermogenic organs, and aerobic capacity

We observed no sexual dimorphism in maximum thermoregulatory metabolic capacity (M_sum_; ANOVA, F = 0.202, p > 0.1). Maximum thermogenic capacity (M_sum_) ranged from 1.373 to 3.309 mL CO_2_/min, with mean = 2.341 mL CO_2_/min ± 0.384 SD. M_sum_ was significantly positively related with T_min_ (GLM, t = 2.764, p < 0.01) and M_b-scaled_ (GLM, t = 2.703, p < 0.01) but not with NDVI (GLM, t = 1.044, p > 0.1).

When restricting the analysis to regional-level, we found that M_sum_ increased alongside with increasing T_min_ (i.e. summer) in *Coastal* (GLM, t = 2.903, p = 0.01) and *Central* (GLM, t = 2.577, p = 0.02) populations alongside increasing T_min_ (i.e. summer), whereas birds from *Inland* showed no significant changes (GLM, t = 1.446, p = 0.163; Figure2E).

At maximum thermogenic capacity, scrub-robins defended similar body temperatures regardless of local T_min_ (GLM; t = 1.510, p > 0.1). Nevertheless, the temperature that triggered the maximum heat production **-** cold limit (Tc_L_) - decreased at lower T_min_ (GLM, t = 7.661, p < 0.01; Figure 2F). This association that was significant for the three regions: *Coastal* (GLM, t = 2.523, p = 0.019), *Central* (GLM, t = 2.634, p = 0.017) and *Inland* (GLM, t = 5.034, p < 0.01).

Overall, the rate of metabolic heat loss was positively associated with increasing T_min_ (GLM; t = 4.858, p < 0.01) but not with M_b-scaled_ (GLM; t = 1.471, p > 0.1). Within-region, thermal conductance significantly increased with increasing T_min_ in *Coastal* and *Central* populations (GLM; t_*Coastal*_= 3.966, p < 0.01; t_*Central*_= 4.976, p < 0.01), while no association with T_min_ was found for *Inland* birds (GLM; t_*Inland*_= 1.962, p = 0.064).

We found M_sum_ to be associated with increasing T_min_ but not NDVI (Table 1, model A). At the whole-organism level, M_sum_ significantly increased with M_b-scaled_ (Table 1, model B), and decreased with fat depots (Table 1, model B). At muscular level, M_sum_ was not affected by mass of thermogenic muscles (Table 1, model C). At the cellular level, none of the proxies of aerobic capacity was associated with M_sum_ (Table 1, model D).

## DISCUSSION

Cold temperatures have long been recognized as an inescapable physiological stressor for small-bodied endotherms such as birds (Tattersall *et al.* 2012). While small birds from north-temperate habitats deal with this challenge through flexibility in thermoregulatory physiological traits (e.g.: (Dawson and Olson 2003), (Vézina *et al.* 2006)), there is a paucity of information on whether sub-tropical species use similar mechanisms (Smit and McKechnie 2010), as well as whether intra-specific variation allows populations to acclimatize to local conditions (van de Ven et al. 2013).

By combining evidence from different levels of organization from populations of a small passerine living in different temperature and primary productivity conditions we found that: i) body condition increased with lower minimum temperatures (T_min_); ii) thermogenic mass remained unchanged regardless of T_min_ or variation in primary productivity (NDVI); iii) in pectoralis muscle, density of mitochondria and lipid droplets was maintained seasonally; iv) in limb muscles, both the mitochondrial and lipid droplet densities decreased with lower T_min_; v) at maximum thermogenic capacity, the environmental temperature that triggered maximum heat production **-** cold limit (Tc_L_) - decreased at lower T_min_ for the three populations; nevertheless, the rate of heat loss remained unchanged for birds experiencing the wider thermal fluctuations - *Inland* birds; vi) maximum thermogenic capacity (M_sum_) showed no seasonal flexibility in the population living in the coldest and least productive area (*Inland*: T_min_ < 0 °C and NDVI = 0.30), and surprisingly increased, in summer, in populations experiencing milder conditions (*Coastal* and *Central* regions) and, vii) M_sum_ was not associated with aerobic capacity in the small limb muscles or in large thermogenic pectoralis.

Overall, these results indicate that at least this arid-zone sub-tropical small bird species does not conform to the cold-adaptation hypothesis or to the climate variability hypothesis. Specifically, the birds that concomitantly experience the coldest conditions and the widest thermal fluctuations showed the least degree of phenotypic flexibility in all eco-physiological traits quantified. Thus, our findings suggest that high M_sum_ and reduced thermal conductance is not the critical mechanism for successful overwintering in the cold winters of the arid sub-tropical habitats.

### Thermogenesis in sub-tropical arid-zones

Although M_sum_ revealed to be seasonally and regionally flexible, the observed patterns contrast with our predictions. Firstly, M_sum_ flexibility was only found in populations living in the milder portion of the scrub-robin range (*Coastal* and *Central)* and not in the population exposed to sub-zero T_min_ in winter (*Inland).* And secondly, the trend was to increase thermogenic capacity in summer, the period where M_sum_ was expected to be at an annual minimum (Swanson and Garland 2009) due to annual highest T_min_ . Our estimates of M_sum_ (assuming an respiratory exchange ratio of 0.71) are, on average, 75% of the values predicted by Swanson and Bozinovic (2011) for a 19 g oscines passerine, suggesting that M_sum_ is not directly linked to cold tolerance in this arid-zone passerine.

Our findings contrast with cold-induced increased of M_sum_ in north-temperate passerines (e.g.: American Goldfinch *Spinus tristis* (Swanson et al. 2014b), House sparrow *Passer domesticus* (Liknes and Swanson 2011a) and are only comparable to results reported for two other southern African birds living in similar latitudes in South Africa: the red-bishop (*Euplectes orix*; (van de Ven *et al.* 2013) and the white-browed sparrow-weaver (*Plocepasser mahali;* (Noakes *et al.* 2017). Both studies found population-specific patterns in M_sum_ flexibility, with some populations increasing M_sum_ in winter (in colder sites), while others showed no seasonal change.

Although the lack of increased M_sum_ in scrub-robins at colder temperatures was unexpected, it was consistent with the unchanged mass of the major thermogenic muscles (pectoralis and gastrocnemius). In contrast to northern-temperate Passeriformes that increase the mass of metabolically active organs (Liknes and Swanson 2011b), we did not observe a cold-induced hypertrophy of pectoralis or gastrocnemius in scrub-robins. Therefore, we postulate that the low productivity in the arid-zone of southern Africa compared to the northern-temperate regions may preclude the growth of the energetically expensive thermogenic muscles. Equally unforeseen was the increased M_sum_ in summer in *Coastal* and *Central* populations. We contend the summer increase in M_sum_ is not related to reproduction as these populations breed late during the Austral winter/spring following predictable rainfall events. Further, the elevated M_sum_ was not related to moulting as we eliminated individuals showing clear signs of moult from analyses. The fact that elevated M_sum_ during summer in *Coastal* populations coincided with elevated heat loss rates, suggest it may be be explained by the strong summer winds at air temperatures below 20°C, typical of the western coast of south Africa (Kruger *et al.* 2010). The exposure to high wind speeds can result in elevated heat loss and hence favour higher thermogenic capacities despite higher T_min_ = 13 °C.

Stressors such as exercise, cold exposure and starvation act to trigger mitochondrial biogenesis (O’Brien 2011) (Putti *et al.* 2015). Thus, we tested the role of density of mitochondria and lipid droplets (energy substrate) as a possible mechanism to respond to the extreme aerobic demand of shivering in the scrub-robin. Our prediction that mitochondrial density in pectoral muscle should increase with lower T_min_, as a mean to power shivering, was not supported (Figure 3B). Yet, the lack of difference in mitochondrial density between *Coastal* and *Inland* birds is reconciled with the lack of flexibility in M_sum_ during winter for the same regions. The significant reduction of lipid droplets in limb in the *Inland* population (T_min_ = -0.4 °C) together with increased number of transversal connections between mitochondria (increases energetic performance (Nasrallah and Horvath 2014)), suggest a depletion of energetic reserves. However, because it did not reflect an increase in thermogenic capacity, we suspect the lipid reserves may have been used to power walk in search for food, in the sparse arid environment.

### Thermosensation prior to thermogenesis

Besides shivering, other mechanisms are proposed to contribute to cold conditions’ tolerance: insulation capacity, locomotor activity and nonshivering thermogenesis. Modulation of insulation from the environment, measured here as flexibility in heat loss at maximum thermogenic capacity (a proxy for thermal conductance), could potentially alter bird’s capacities to better control heat loss. However, our estimates, besides showing no clear regional differences in termal conductance, also revealed no seasonal differences in thermal conductance in the population experiencing the coldest temperatures (*Inland*).

Increasing locomotor activity exploits the thermodynamic inefficiency of catabolic reactions in skeletal muscles (60% energy released as heat) and nonshivering thermogenesis uses a deflection of proton flux in mitochondria from ATP production to heat production (Hochachka and Somero 2002). However, none of the mechanisms seem plausible in the scrub-robin. First, because being on the move for the sake of heat production in low productivity environments may lead to a mismatch between energetic demand and supply. And secondly, recent work suggests that nonshivering thermogenesis is not a prominent contributor to thermogenesis in adult passerines (Cheviron and Swanson 2017). With neither locomotor activity nor nonshivering thermogenesis responsible for thermoregulation, we propose the explanation may be on how the scrub-robins sense the cold. This ideas is supported by the fact we recorded a drastic reduction in T_b_ to 34 – 36 °C, while birds maintained their maximum metabolic capacity, and remarkably that scrub-robins could fly at ease at T_b_ = 30 - 34 °C, soon after the cold-experiment. It is thus possible that intra-specific variation exists in the somatosensory system (cutaneous thermoreceptors; e.g.: TPRM8 channel (Matos-Cruz *et al.* 2017)) and hence some populations tolerated prolonged exposure to cold and hypothermia without engaging in active production of heat. An understanding of the sensory perception of cold in Karoo scrub-robin, as the first system to mediate organism-environment interactions (Gracheva and Bagriantsev 2015), is certainly warranted.

To conclude, we believe our study exemplifies that only when implementing a detailed framework at the intra-specific level one can contemplate to understand the factors underlying phenotypic flexibility and ultimately its role in adaptation.

## Acknowledgments

We are grateful to Hilda Vermuelen, Elsa van Schalkwyk and family, Eugene Marinus and Colleen Rust (SANBI Botanical Garden in Nieuwoudtville), and Marelé Nel for their support during field expeditions. We thank the South African Provincial Authorities for providing permits. This study was supported by a Marie Sklodowska-Curie Individual fellowship (European Union Horizon 2020 Research and Innovation Programme, grant 655150 - BARREN) and a British Ornithologists’ Union small grant to AMR.

## Author contributions

AMR and BS conceived the study. AMR collected field data and conducted statistical analyses with critical input from CP, MTPG and BS. AMR and CP performed immunostaining and confocal microscopy imaging. NBP and BS collected physiological data. All authors wrote the manuscript and gave final approval for publication.

## Data accessibility

Data used in this study is provided in a spreadsheet as Supplementary Information.

